# Identification of the C9-Hydrogenase for 9,17-Dioxo-1,2,3,4,10,19-Hexanorandrostan-5-oic Acid (9,17-DOHNA) and the 7α-Dehydratase Essential for Initiating β-Oxidation of the B-, C-, and D-Rings in Steroid Degradation by *Comamonas testosteroni* TA441

**DOI:** 10.1101/2025.11.20.689525

**Authors:** Masae Horinouchi

## Abstract

*Comamonas testosteroni* TA441 is a model aerobic steroid-degrading bacterium whose sterane degradation pathway has been elucidated in the greatest detail to date. Similar pathways have been identified in many genera of bacteria, including both proteobacteria and actinobacteria such as *Mycobacterium tuberculosis*. However, the genes encoding the C9-hydrogenase for 9,17-dioxo-1,2,3,4,10,19-hexanorandrostan-5-oic acid (9,17-DOHNA, also known as HIP) and the 7α-dehydratase essential for initiating β-oxidation of the B-, C-, and D-rings had not been identified within the gene cluster responsible for B-, C-, and D-ring degradation.

In this study, we identified these missing genes, located adjacent to the *chsE1E2H1H2ltp2* cluster involved in C17 side-chain degradation, and designated them *scdB* and *scdH*, respectively. This finding completes the elucidation of all degradation steps of 9,17-DOHNA prior to D-ring cleavage. AlphaFold-predicted models showed that the hydrogenases/dehydrogenases involved in steroid degradation in TA441—ScdG, ScdE, 3β-dehydrogenase (3β-DH), SteA, 3α-dehydrogenase (3α-DH), and SteB—share a similar Rossmann-like α/β/α sandwich fold with ScdB and function as dimers. In contrast, ScdH was predicted to form a homohexameric structure, similar to ScdY and ScdN, members of the crotonase-like enoyl-CoA hydratase/isomerase family involved in B-, C-, and D-ring degradation. Furthermore, AlphaFold modeling revealed that SteC, the dehydratase responsible for removing the C12β-hydroxyl group from 9,17-DOHNA derivatives, exhibits strong structural similarity to BaiE, the bile acid 7α-dehydratase of *Clostridium scindens* JCM 10418/VPI 12708, despite sharing only ∼28% amino acid sequence identity.

## IMPORTANCE

Research on bacterial aerobic steroid degradation began more than 70 years ago, initially to produce intermediates for steroid drug synthesis. Recently, this field has gained renewed attention due to its implications for human health—for example, the role of cholesterol import and degradation in the persistence of *Mycobacterium tuberculosis* H37Rv within chronically infected lungs.

*Comamonas testosteroni* TA441 serves as a key model organism for elucidating aerobic steroid degradation, with pathways for cleavage of the A-, B-, C-, and D-rings already well established. The functions and structures of the enzymes identified in TA441 display striking similarities to those in actinobacteria such as *M. tuberculosis*.

In this study, we identified two enzymes indispensable for initiating β-oxidation of the B-, C-, and D-rings, thereby filling the last remaining gaps for initiationg this pathway. Our AlphaFold-based structural analysis of these enzymes not only provides new insights into the steroid metabolism of *M. tuberculosis* but also broadens understanding of the ecological and physiological significance of bacterial steroid degradation.

## INTRODUCTION

Steroid compounds perform a wide range of biological functions in plants and animals, including humans, where they serve essential roles as hormones, cholesterol, and bile acids (1–4). Aerobic degradation of steroids by bacteria has been recognized for over 70 years, with pioneering studies in the 1950s and 1960s using the two representative steroid-degrading bacteria, the actinomycete *Rhodococcus equi* and the pseudomonad *Comamonas testosteroni* (5–8).

Our previous studies elucidated the almost complete degradation pathways of the sterane structure—A-, B-, C-, and D-rings—as well as several side chains such as hydroxyl substituents and the C17 isopropyl residue in *C. testosteroni* TA441, establishing this strain as a model organism for aerobic bacterial steroid degradation (9–29). (The whole degradation pathway revealed in TA441 is shown in Supplementary Materials Fig. S1). Genes for A- and B-ring cleavage via A-ring aromatization and those for B-, C-, and D-ring degradation primarily via β-oxidation are located at both ends of the 120-kb “mega-cluster” of steroid degradation genes (Fig. 1).

**Fig 1.**
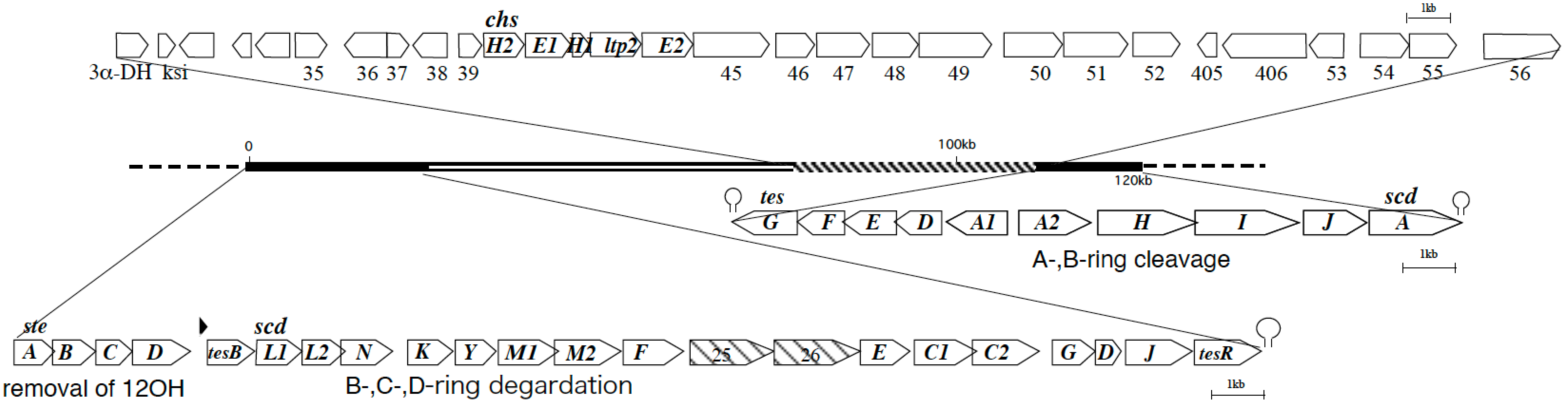
The 120-kb mega-cluster of steroid degradation genes in *C. testosteroni* TA441. The *tesG3scdA* cluster contains genes mainly responsible for aromatization of the A-ring (*tesH, tesI, tesJ*) and for ring cleavage and degradation of the A- and B-rings (*tesG, tesF, tesE, tesD, tesA1A2*). The *steA3tesR* cluster contains genes required for removal of the C12³ hydroxyl group (*steABCD*) and genes involved in concurrent cleavage and ³-oxidation of the C- and D-rings (*scdL1L2, scdN, scdK, scdY, scdM1M2, scdF, scdE, scdC1C2, scdG, scdD, scdJ*). *tesB* encodes the meta-cleavage enzyme for the aromatized A-ring, and *tesR* encodes the positive regulator of steroid degradation genes. *scdA* encodes the CoA-transferase initiating ³-oxidation of the B-, C-, and D-rings. Other genes: 3³-DH, encoding 3³-hydroxydehydrogenase; *ksi*, encoding 3-ketosteroid —435 isomerase; *chsH2, chsE1, chsH1, trp2, chsE2,* which encode enzymes for removal of the isopropyl residue at C17. Genes between ORF35355 are predicted to be steroid-degradation genes yet to be characterized.

In bacterial cholic acid degradation, the C17 side chain must be shortened to a ketone via a propionyl residue prior to sterane-ring degradation. Our most recent work demonstrated that this conversion is mediated by the dehydrogenase ChsE1E2, hydratase ChsH1H2, and lipid transfer protein Ltp2, which share high structural similarity with the corresponding enzymes in *Mycobacterium tuberculosis* (28, 30–32). Aerobic steroid degradation pathways similar to that of TA441 are thought to exist in other aerobic steroid-degrading bacteria, including actinomycetes such as *Rhodococcus*, *Mycobacterium*, and *Sphingomonas* (33–36), as well as pseudomonadota such as *Pseudomonas* spp. (37), *Steroidobacter denitrificans* (38), *Caenibius tardaugens* (39), and other Comamonadaceae. In *M. tuberculosis* H37Rv, whose cholesterol import system is essential for persistence in chronically infected animal lungs (40), cholesterol catabolism is also critical for pathogenic maintenance in the host (41).

Among at least 27 recognized *Comamonas* species, only *C. aquatica*, *C. kerstersii*, *C. terrigena*, and *C. testosteroni* are considered opportunistic pathogens, and only *C. thiooxydans* (42) and *C. resistens* (43, 44) have been reported to possess steroid degradation genes similar to those of *C. testosteroni*.

During A-ring cleavage, aromatization of the A-ring and opening of the B-ring generate intermediates that undergo degradation analogous to the bacterial biphenyl degradation pathway, forming 9,17-dioxo-1,2,3,4,10,19-hexanorandrostan-5-oic acid (9,17-DOHNA; also known as HIP, 3aα-H-4α[3′-propionic acid]-7aβ-methylhexahydro-1,5-indanedione) and 2-hydroxyhexa-2,4-dienoic acid (Fig. 2 and Fig. S1) (9–16, 22, 26). The intermediate 9,17-DOHNA, containing the intact C-and D-rings and an opened B-ring, is converted into its CoA ester by ScdA, encoded in the A/B-ring cleavage cluster, and subsequently degraded by successive β-oxidation cycles mediated by Scd genes located in the B,C,D-ring degradation cluster (Fig. 1) (17–21, 23–26, 29).

**Fig. 2.**
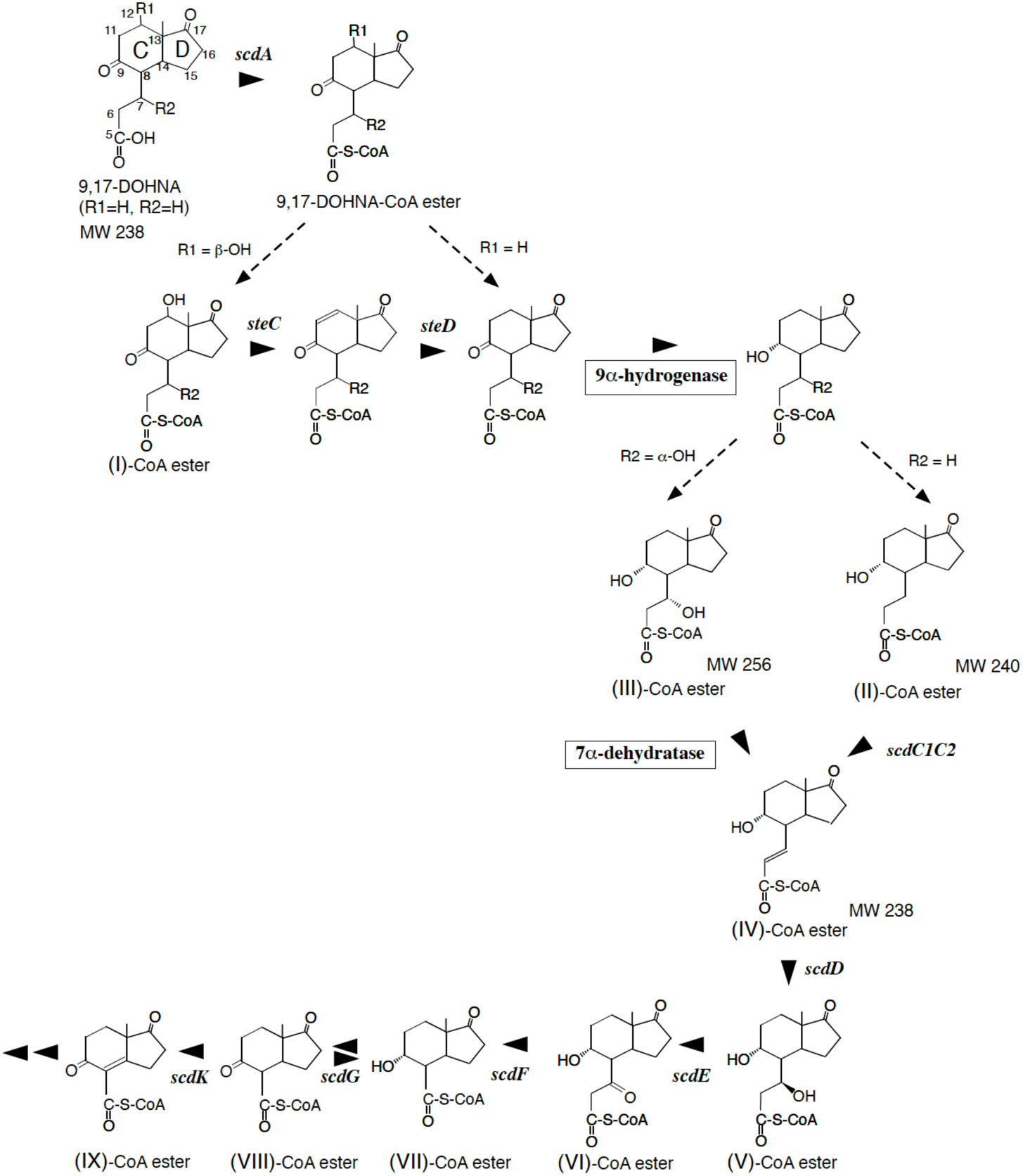
Steroid degradation pathway of the B-, C-, and D-rings, from 9,17-dioxo-1,2,3,4,10,19-hexanorandrostan-5-oic acid (9,17-DOHNA) to **IX**-CoA ester, immediately before D-ring cleavage, in *C. testosteroni* TA441. Enzymes responsible for C9³-hydrogenase and C7³-dehydratase activities (boxed) remain unidentified. Compounds are; 9,17-DOHNA; 9,17-dioxo-1,2,3,4,10,19-hexanorandrostan-5-oic acid (also known as 3a³-*H*-4³ {3′-propionic acid}-7a³-methylhexahydro-1,5-indanedione, HIP) (R1, R2=H), (**I**); 12³-hydroxy-9,17-dioxo-1,2,3,4,10,19-hexanorandrostan-5-oic acid (R2=H), (**II**); 9³-hydroxy-17-oxo-1,2,3,4,10,19-hexanorandrostan-5-oic acid, (**III**); 7³,9³-dihydroxy-17-oxo-1,2,3,4,10,19-hexanorandrostan-5-oic acid, (**IV**); 9³-hydroxy-17-oxo-1,2,3,4,10,19-hexanorandrost-6-en-5-oic acid, (**V**); 7³,9³-dihydroxy-17-oxo-1,2,3,4,10,19-hexanorandrostan-5-oic acid, (**VI**); 9³-hydroxy-7,17-dioxo-1,2,3,4,10,19- hexanorandrostan-5-oic acid, (**VII**); 9³-hydroxy-17-oxo-1,2,3,4,5,6,10,19-octanorandrostan-7-oic acid, (**VIII**); 9,17-dioxo-1,2,3,4,5,6,10,19-octanorandrostan-7-oic acid, and (**IX**); 9,17-dioxo-1,2,3,4,5,6,10,19-octanorandrost-8(14)-en-7-oic acid. Enzymes are; SteC (dehydradase for 12³-OH to produce a double at C10(12)), SteD (reductase for a double at C10(12) to a single bond), ScdA (CoA-transferase for 9,17-DOHNA), ScdC1C2 (—6-dehydrogenase for **II-**CoA ester), ScdD (**IV-**CoA ester —6-hydratase), ScdE (**V-**CoA ester dehydrogenase at C7), ScdF (**VI-**CoA ester thiolase/CoA-transferase), ScdG (hydrogenase primarily for 9-OH of **VII-**CoA ester), and ScdK (—8(14)-dehydrogenase for **VIII-**CoA ester).

Although the overall steroid degradation pathway in TA441 is well characterized, several expected enzymatic steps have remained unidentified. Here, we report the discovery of the C9-hydrogenase acting on 9,17-DOHNA-CoA and the 7α-dehydratase essential for initiating β-oxidation of the B,C,D-rings in steroid degradation by *C. testosteroni* TA441. We also present AlphaFold-based analyses of these enzymes together with previously characterized steroid-degrading enzymes of TA441.

## RESULTS

### Analysis of the culture of ORF38- and ORF39-disrupted mutants incubated with cholic acid and its analogs

*Comamonas testosteroni* TA441 degrades steroids through C17-side chain degradation (28), A,B-ring cleavage, which is analogous to the bacterial “meta-cleavage pathway” involved in the degradation of aromatic compounds such as biphenyl (9–16), followed by B-, C-, D-ring degradation via several cycles of β-oxidation (17–21). The addition of CoA by ScdA to the C,D-ring–containing intermediate with a cleaved B-ring, 9,17-dioxo-1,2,3,4,10,19-hexanorandrostan-5-oic acid (9,17-DOHNA), also known by its IUPAC name 3aα-H-4α[3′-propionic acid]-7aβ-methylhexahydro-1,5-indanedione (HIP), initiates the B,C,D-ring degradation (Fig. 2). This step is followed by 9α-hydrogenation, which produces

9α-hydroxy-17-oxo-1,2,3,4,10,19-hexanorandrostan-5-oic acid (**II**). Removal of the 12β-hydroxyl group by SteC and SteD occurs prior to the 9α-hydrogenation (Fig. 2), as mutants disrupted in *scdC* (SteC^−^) and *scdD* (SteD^−^) accumulate compounds containing a ketone group at C9 (23). The 9α-hydrogenase acting on 9,17-DOHNA-CoA ester has not been identified; although ScdG exhibits 9α-hydrogenation/dehydrogenation activity, it is not the primary enzyme responsible for this step because the ScdG-disrupted mutant retains normal 9α-hydrogenation activity toward 9,17-DOHNA-CoA ester. After the 9α-hydrogenation step, a double bond is introduced by ScdC1C2 when the initial steroid lacks a hydroxyl group at C7, whereas a C7α-dehydratase introduces the double bond when the steroid contains an α-oriented hydroxyl group at C7 (Fig. 2) (26).

C7α-dehydration was suggested to be reversible, as 7α,9α-dihydroxy-17-oxo-1,2,3,4,10,19-hexanorandrostan-5-oic acid (**III**) was detected in the culture of the *scdD*-disrupted mutant (ScdD^−^) incubated with testosterone (Supplementary Materials Fig. S2). However, the enzyme responsible for C7α-dehydration had not been identified previously.

We analyzed the cultures of ORF38- and ORF39-disrupted mutants (ORF38^−^ and ORF39^−^, respectively) incubated with cholic acid as preliminary experiments. These ORFs are located just upstream of *chsH2E1H1ltp2chsE2*, which encode enzymes involved in degradation of the propionyl residue during C17-side chain breakdown (Fig. 1) (17, 20, 24, 28). The results suggested that ORF38 and ORF39 participate in the early steps of B,C,D-ring degradation. Homology searches indicated that ORF38 and ORF39 encode an enoyl-CoA hydratase and an SDR-family oxidoreductase, respectively.

Therefore, we incubated ORF38^−^, ORF39^−^, and the mutants ScdA^−^, ScdC1C2^−^, and ScdD^−^ (Fig. 2) with cholic acid (CA), deoxycholic acid (DC), chenodeoxycholic acid (CD), and lithocholic acid (LT) for 7 days, followed by UPLC/MS analysis (Fig. 3). The ORF38^−^ mutant accumulated a large amount of a compound with *m/z* 255 at RT = 2.99 min in cultures supplemented with CA and CD, both possessing an α-oriented hydroxyl group at C7, but showed no notable accumulation of intermediate compounds with DC or LT, which lack a hydroxyl group at C7 (Fig. 3A; see also Supplementary Materials Fig. S2). A small amount of **III** was detected in ScdD^−^ cultures, displaying the same RT (2.99 min) as that of the compound in the ORF38^−^ culture, suggesting that this compound corresponds to **III** and that the enzyme encoded by ORF38 is the dehydratase acting on the **III**-CoA ester. 9,17-DOHNA derivatives containing a 7α-hydroxyl and/or 12β-hydroxyl group (with *m/z* 269 or 253) and 9,17-DOHNA (*m/z* 237) were expected to be detected from the ORF38^−^ cultures incubated with CA, DC, CD, and LT, respectively. As shown in Fig. 3B, these compounds were indeed detected: the derivatives with 7α- and 12β-hydroxyl groups (*m/z* 269, RT = 0.61 min) and those with only the 7α-hydroxyl group (*m/z* 253, RT = 1.77 min) appeared predominantly as parent ions with *m/z* 207 and 191, respectively (Fig. 3B3).

**Fig. 3.**
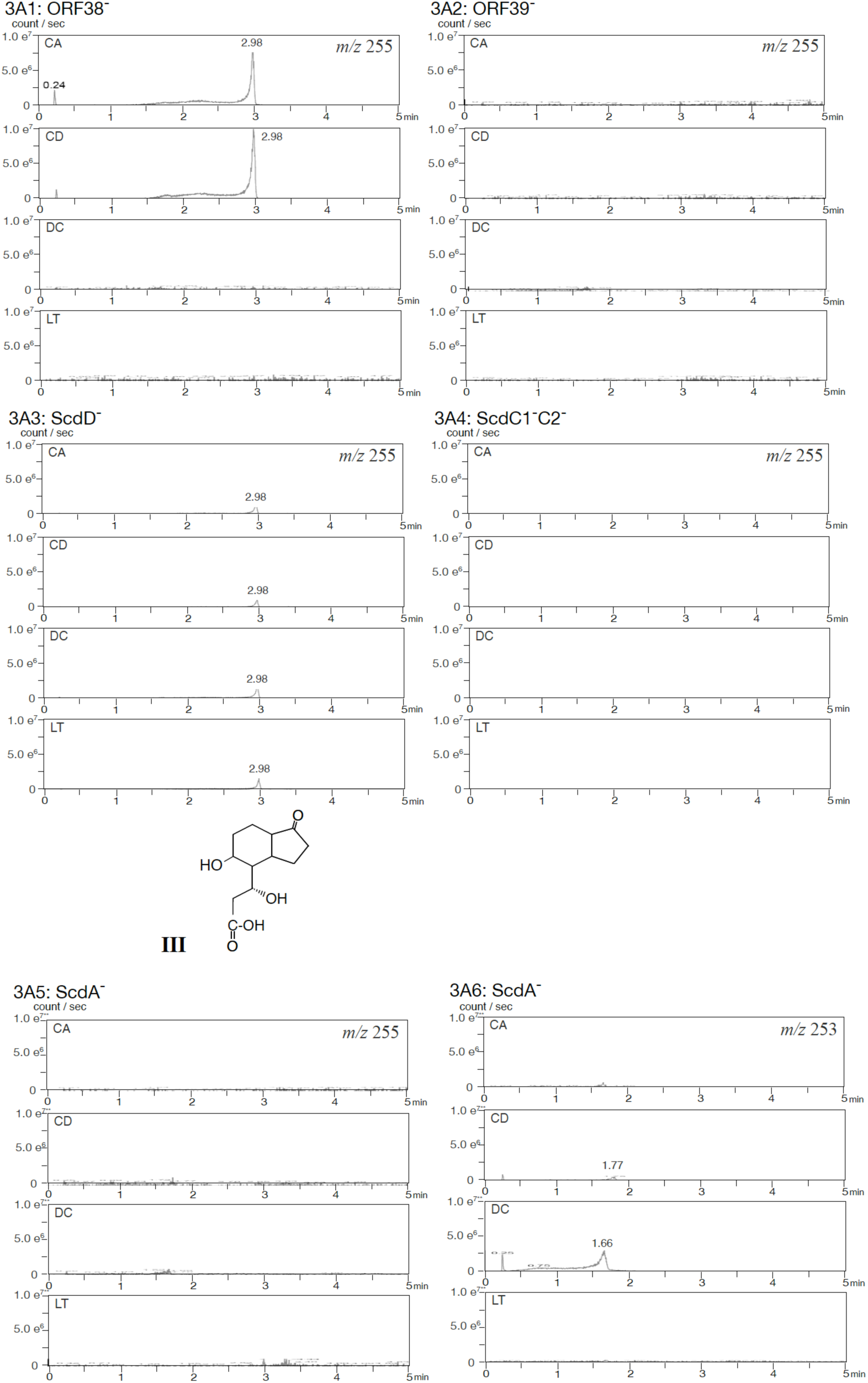
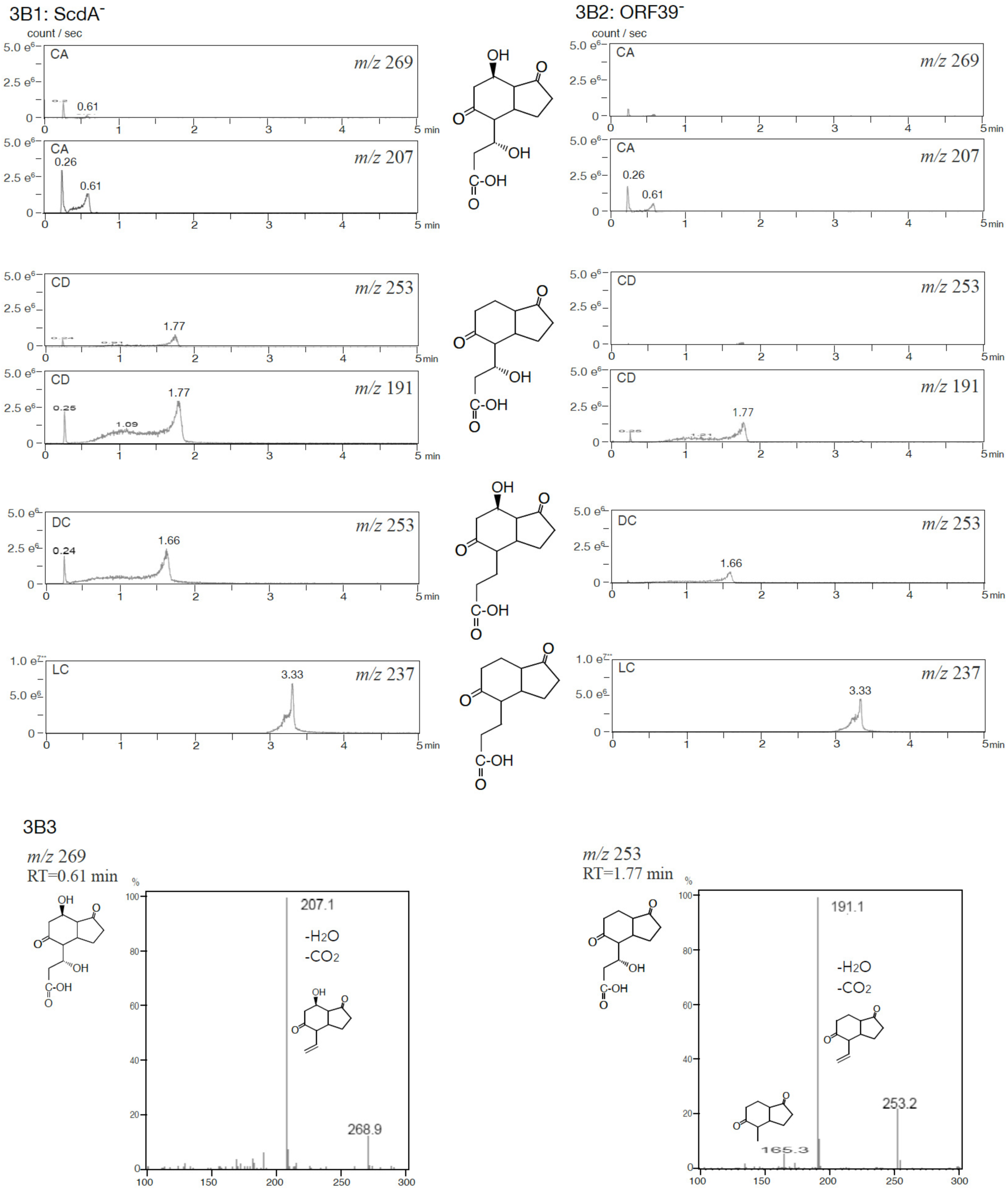
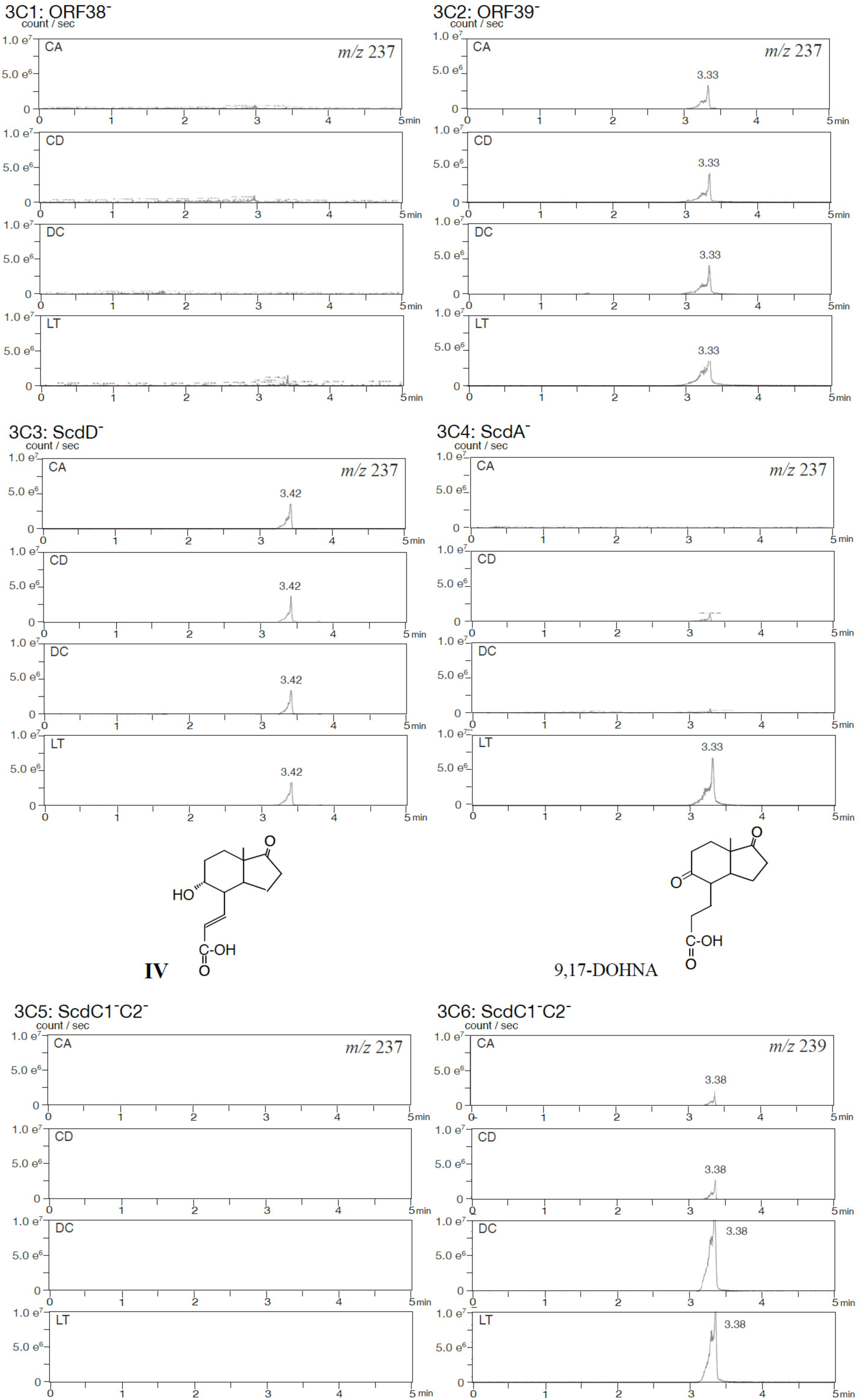
UPLC/MS analysis of ORF38^3^ and ORF39^3^ mutants incubated with 0.05% cholic acid (CA), chenodeoxycholic acid (CD), deoxycholic acid (DC), or lithocholic acid (LT) for 7 d (3A1, 3A2, 3B1, 3B2). Mutants ScdD^3^ (3A3), ScdC1^3^C2^3^ (3A4), and ScdA^3^ (3A5, 3A6) were analyzed similarly for peak identification. Mutants were constructed by inserting a kanamycin resistance gene. Peaks in mass spectra represent: *m/z* 255 at RT = 2.89 min (7³,9³-dihydroxy-17-oxo-1,2,3,4,10,19-hexanorandrostan-5-oic acid, **III**) (3A1CA, 3A1CD, and 3A3); *m/z* 253 at RT = 1.77 min (7³-hydroxy-9,17-dioxo-1,2,3,4,10,19-hexanorandrostan-5-oic acid) (3A6 CD, 3B1CD, and 3B2 CD with mass spectrum in 3B3); *m/z* 253 at RT = 1.66 min (12³-hydroxy-9,17-dioxo-1,2,3,4,10,19-hexanorandrostan-5-oic acid) (3A6 DC, 3B1DC, and 3B2 DC); *m/z* 269 at RT = 0.61 min (7³,12³-dihydroxy-9,17-dioxo-1,2,3,4,10,19-hexanorandrostan-5-oic acid) (3B1CA and 3B2CA with mass spectrum in 3B3); *m/z* 237 at RT = 3.33 min (9,17-dioxo-1,2,3,4,10,19-hexanorandrostan-5-oic acid, 9,17-DOHNA)(3B1LT, 3B2LT, 3C2, and 3C4LT); *m/z* 237 at RT = 3.42 min (9³-hydroxy-17-oxo-1,2,3,4,10,19-hexanorandrost-6-en-5-oic acid, **IV**) (3C3); and *m/z* 239 at RT = 3.38 min (9³-hydroxy-17-oxo-1,2,3,4,10,19-hexanorandrostan-5-oic acid, **II**) (3C6). The vertical axis indicates intensity (counts/sec), and the horizontal axis indicates retention time (min).

The ORF39^−^ mutant accumulated a compound with *m/z* 237 at RT = 3.33 min in all cultures containing cholic acid derivatives (Fig. 3C). Among the intermediate compounds shown in Fig. 2, both 9,17-DOHNA and 9α-hydroxy-17-oxo-1,2,3,4,10,19-hexanorandrost-6-en-5-oic acid (**IV**) have a molecular weight of 238. 9,17-DOHNA accumulates primarily in the ScdA^−^ mutant, while **IV** accumulates in the ScdD^−^ mutant, with the latter producing a distinct peak at *m/z* 175 (Fig. 3C; Supplementary Materials Fig. S3). Therefore, the compound with *m/z* 237 at RT = 3.33 min in the ORF39^−^ culture is identified as 9,17-DOHNA, indicating that ORF39 encodes the 9α-hydrogenase responsible for the conversion of 9,17-DOHNA-CoA ester.

### Complementation experiments with ORF38 and ORF39 disrupted mutants

For further confirmation, the broad-host-range plasmid pMFYMhpRA (derived from pMFY42) (24, 45), serving as a negative control, and the same vector carrying either ORF38 (pMFYMhpORF38) or ORF39 (pMFYMhpORF39) were introduced into the ORF38^−^ and ORF39^−^ mutants, respectively (Table 1 and Supplementary Materials Table S1). pMFYMhpRA was designed to express cloned genes upon addition of 3-(3-hydroxyphenyl)propionic acid (3HPP), since in previous studies on sterane ring degradation, only partial activity recovery, sometimes hardly detectable recovery, was observed when the substrate and product of the target enzyme were CoA esters.

**Table 1.**
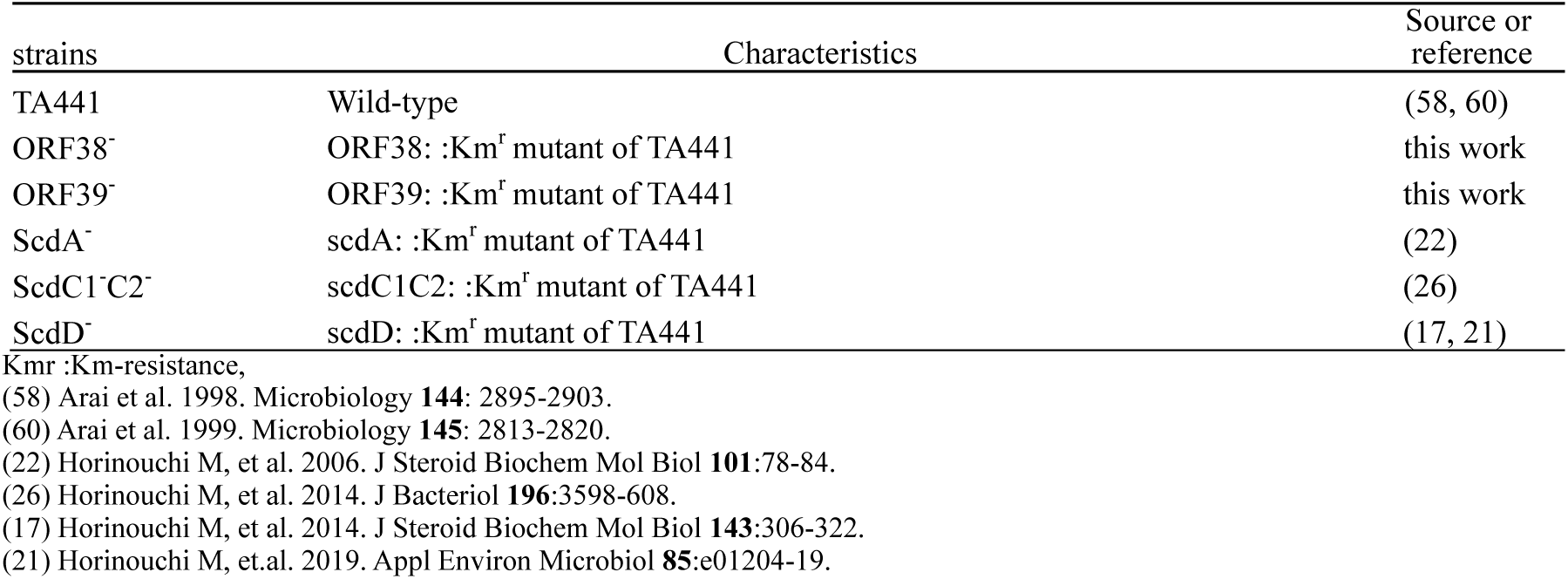
strains.

| strains | Characteristics | Source or reference |
| --- | --- | --- |
| TA441 | Wild-type | (58, 60) |
| ORF38<sup>-</sup> | ORF38: :Km<sup>r</sup> mutant of TA441 | this work |
| ORF39<sup>-</sup> | ORF39: :Km<sup>r</sup> mutant of TA441 | this work |
| ScdA<sup>-</sup> | scdA: :Km<sup>r</sup> mutant of TA441 | (22) |
| ScdC1C2<sup>-</sup> | scdC1C2: :Km<sup>r</sup> mutant of TA441 | (26) |
| ScdD<sup>-</sup> | scdD: :Km<sup>r</sup> mutant of TA441 | (17, 21) |

Kmr :Km-resistance,
(58) Arai et al. 1998. Microbiology **144**: 2895-2903.
(60) Arai et al. 1999. Microbiology **145**: 2813-2820.
(22) Horinouchi M, et al. 2006. J Steroid Biochem Mol Biol **101**:78-84.
(26) Horinouchi M, et al. 2014. J Bacteriol **196**:3598-608.
(17) Horinouchi M, et al. 2014. J Steroid Biochem Mol Biol **143**:306-322.
(21) Horinouchi M, et.al. 2019. Appl Environ Microbiol **85**:e01204-19.

The ORF38^−^ and ORF39^−^ mutants were cultured with CD and LT, respectively, for 1 day before adding 3HPP, and the cultures were analyzed by UPLC/MS at appropriate time intervals. Figure 4A shows chromatograms of *m/z* 255 (**III**) and *m/z* 237 (**IV**) from cultures of ORF38^−^ carrying either pMFYMhpORF38 or the empty vector pMFYMhpRA. Compound **IV** was detected only in the complemented strain, demonstrating that the enzyme encoded by ORF38 catalyzes dehydration of **III**-CoA ester to **IV**-CoA ester. This enzyme was thus designated ScdH.

**Fig. 4.**
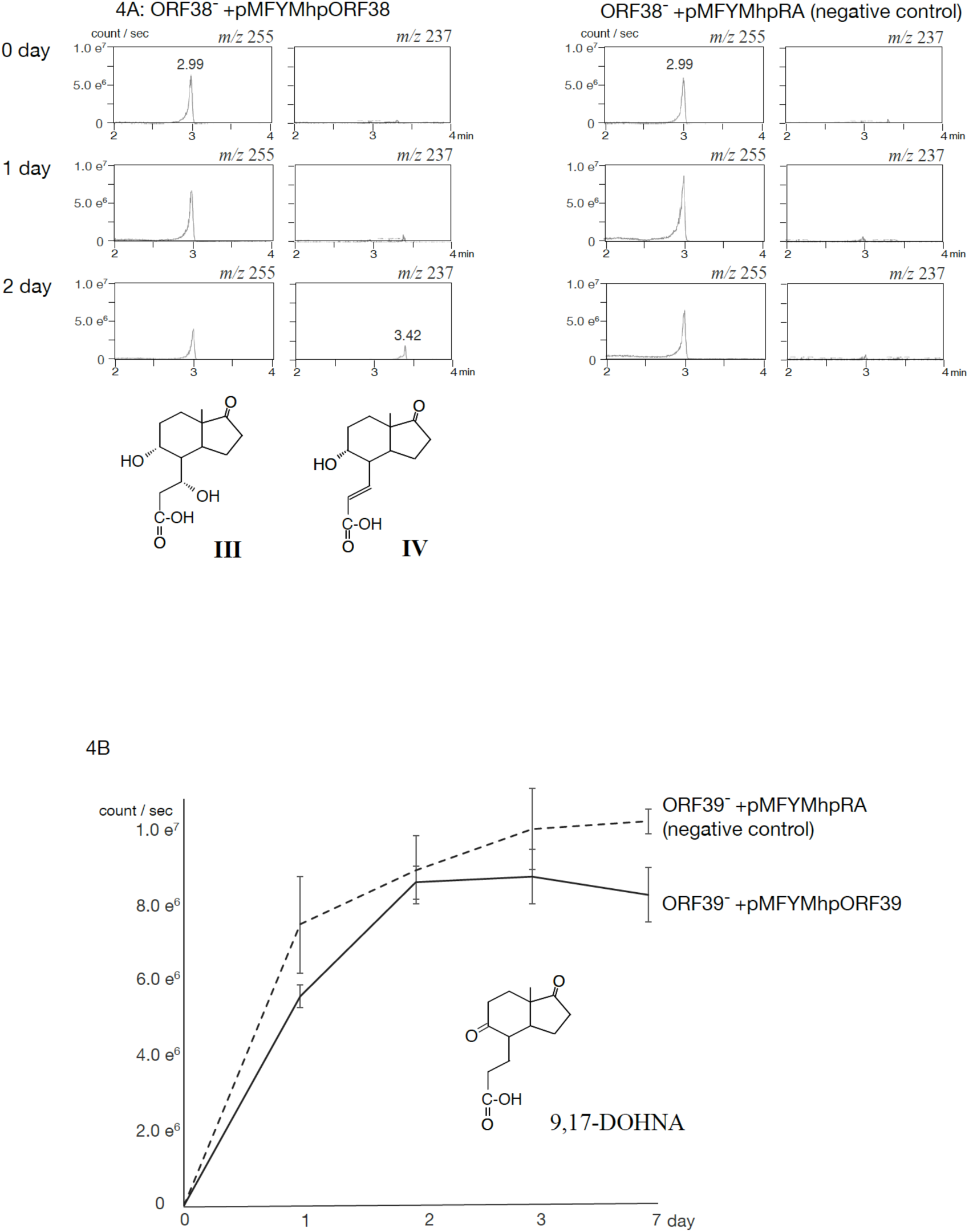
Complementation analysis of ORF38^3^ and ORF39^3^ mutants. **4A**: Mass chromatograms of ORF38^3^ carrying pMFYMhpORF38 and the vector control pMFYMhpRA, incubated with 0.05% LT for 032 d, showing peaks at *m/z* 255 (**III**) and *m/z* 237 (**IV**). **4B**: Amount of 9,17-DOHNA accumulated in ORF39^3^ expressing pMFYMhpORF39 or vector control, incubated with 0.05% 1,4-androstadien-3,17-dion (ADD) for 7 d. Quantification was based on mass chromatograms.

In contrast, chromatograms of ORF39^−^ carrying pMFYMhpORF39 and ORF39^−^carrying the control vector showed only minor differences even after induction with 3HPP. To validate the catalytic activity, we tested the conversion of 9,17-DOHNA-CoA ester using crude extracts of *E. coli* harboring pUCORF39 (a pUC19-based plasmid carrying ORF39, Supplementary Materials Table S1). ORF39^−^ was cultured with 0.1% 1,4-androstadien-3,17-dione (ADD) for 7 days, then sonicated, and the supernatant was used as substrate. ADD was used because it is more readily converted to 9,17-DOHNA by TA441 than LT. *E. coli* strains carrying either pUCORF39 or pUC19 (control) were cultured with ampicillin and IPTG for 1 day, sonicated, and incubated with the substrate solution at 37°C for 1 h. HPLC/MS analysis revealed that the amount of 9,17-DOHNA detected in the reaction with pUCORF39 (1.44 × 10⁷ counts/s, SD = 3.42 × 10⁶) was approximately 15% lower than in the control (1.75 × 10⁷ counts/s, SD = 3.53 × 10⁶). Because the decrease was modest, additional experiments were conducted with ORF39^−^carrying pMFYMhpORF39 and the control strain, using ADD as substrate without tetracycline supplementation, since tetracycline was found to inhibit dehydrogenase ChsE1E2 in previous studies (28). Repeated experiments confirmed that the amount of 9,17-DOHNA was consistently lower in the complemented strain (Fig. 4B). Thus, ORF39 was identified as encoding the 9α-hydrogenase responsible for converting 9,17-DOHNA-CoA ester to **III-**CoA (C7=αOH) and/or **II**-CoA (C7=H), and this enzyme was named ScdB.

During C- and D-ring degradation, ScdG functions as a 9α-dehydrogenase, catalyzing the oxidation of 9α-hydroxy-17-oxo-1,2,3,4,5,6,10,19-octanorandrostan-7-oic acid (**VII**)-CoA ester to 9,17-dioxo-1,2,3,4,5,6,10,19-octanorandrostan-7-oic acid (**VIII**)-CoA ester. The conversion experiments yielded similarly small amounts of product (18), likely because CoA esters possessing a C9-ketone group and a C8–14 single bond are unstable and readily interconvert with compounds containing a C9-ketone and a C8–14 double bond or a C9-hydroxyl and a C8–14 single bond during steroidal C- and D-ring degradation in TA441 (18, 24).

### AlphaFold predicted three-dimensional (3D) structures of ScdB and ScdH; comparison with other hydrogenases and hydratases in steroid degradation

ScdB identified in this study catalyzes the similar reaction, interconversion between a β-oriented hydroxyl group and a ketone group at C9, to previously identified ScdG (Fig. 1) (18). Homology search indicated that both ScdB and ScdG belong to the SDR family of oxidoreductases, therefore, we compared the AlphaFold three-dimensional (3D) models of (ScdB)₂ and (ScdG)₂ (Fig. 5A1 and 5A2; expected position errors and alignments of the five top models are shown in Supplementary Materials Fig. S4). Both ScdB and ScdG were predicted to function as homodimers, and their overall folds were similar, each exhibiting the characteristic Rossmann-like α/β/α sandwich fold typical of SDR oxidoreductases.

**Fig. 5.**
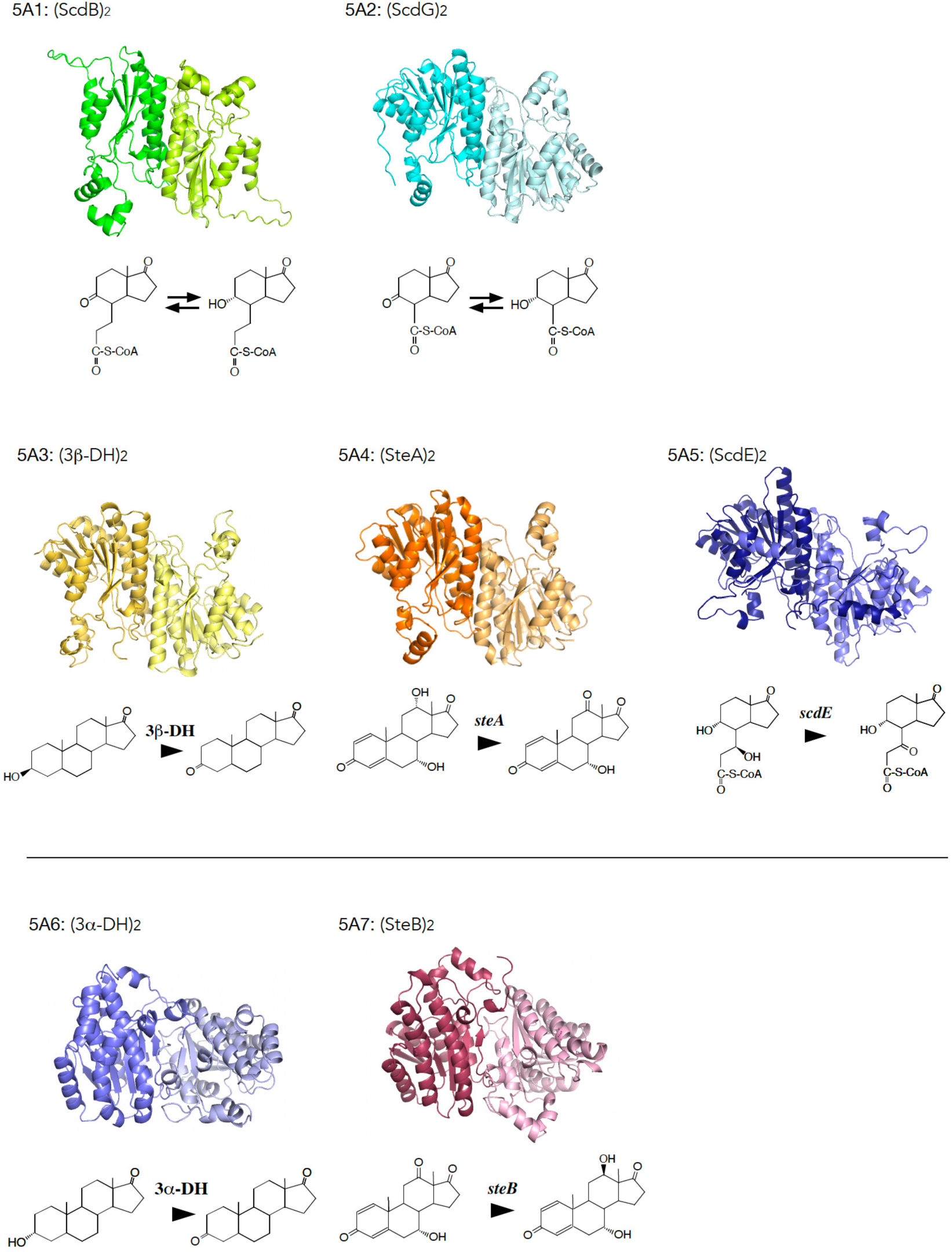
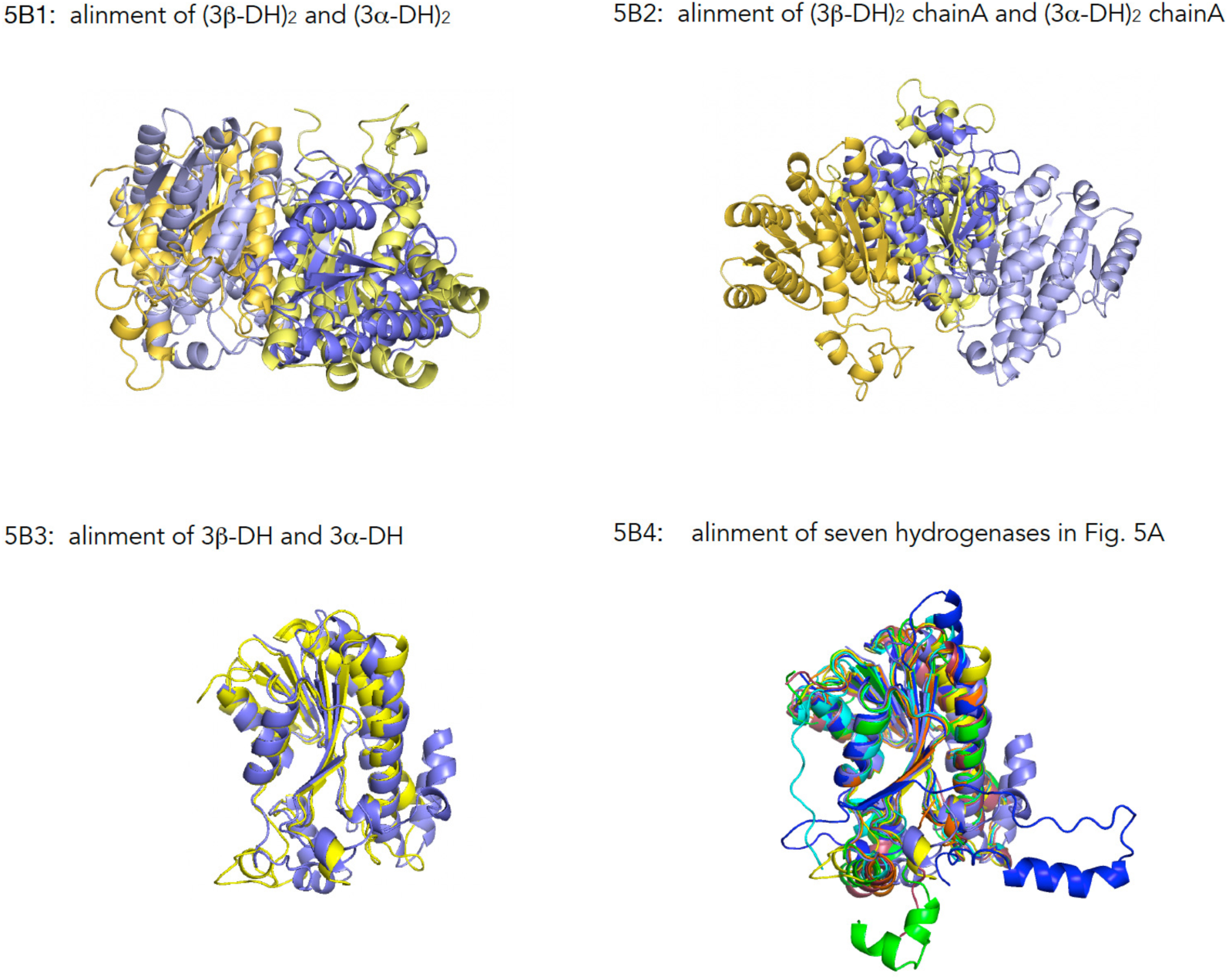
AlphaFold-predicted 3D structures of (ScdB)¢ (5A1) and other hydrogenases/dehydrogenases involved in interconversion of hydroxyl and ketone groups, (ScdG)_2_ (**VII**-CoA ester to **VI**-CoA ester) (5A2), (ScdE)_2_ (**V**-CoA ester to **VI**-CoA ester) (5A3), (3³-DH)_2_ (3³-dehydrogenase for compounds with sterane structure) (5A4), (SteA)_2_ (12³-dehydrogenase) (5A5), (3³-DH)_2_ (3³-dehydrogenase for compounds with sterane structure) (5A6), and (SteB)_2_ (12³-hydrogenase) (5A7). Reactions catalyzed by each enzyme are shown below the structures. Expected position errors and alignments of top models are shown in Supplementary Materials Fig. S4A to S4C. **5B:** Alignment of (3³-DH)¢ and (3³-DH)¢ (5B1), alignment of the monomers (5B2), and monomer-level alignment of all enzymes in Fig. 5A (5B3).

In contrast, the AlphaFold model of (ScdK)₂—an enzyme involved in the downstream reaction after ScdG (Fig. 1) and belonging to the NAD(P)H-dependent flavin oxidoreductase YrpB nitropropane dioxygenase family—was markedly different from those of ScdB and ScdG (Supplementary Materials Fig. S5A1; expected position errors and alignments of the top models in Fig. S5B). In *M. tuberculosis*, the enzymes corresponding to ScdG and ScdK were reported to function in the opposite order, catalyzing C8–14 dehydrogenation and 9α-hydrogenation/dehydrogenation, respectively (34). The AlphaFold model of only SteD (C11 alkene reductase; Fig. 1) resembled ScdK overall among hydrogenases/dehydrogenases for steroid degradation identified to date in TA441, comprising eight α-helices forming a ring-like structure (Supplementary Materials Fig. S5A2,B), but alignment of ScdK and SteD was unsuccessful.

In addition to (ScdB)₂ and (ScdG)₂, we predicted the 3D structures of all hydrogenases/dehydrogenases identified to date in TA441 that catalyze interconversion between a ketone group and a hydroxyl group during steroid degradation. These included (3β-DH)₂ (3β-dehydrogenase for steranes), (SteA)₂ (12α-dehydrogenase for steranes), (3α-DH)₂ (3α-dehydrogenase for steranes), (SteB)₂ (12β-hydrogenase for steranes), and (ScdE)₂ (responsible for converting 7β,9α-dihydroxy-17-oxo-1,2,3,4,10,19-hexanorandrostan-5-oic acid (**V**)–CoA ester to 9α-hydroxy-7,17-dioxo-1,2,3,4,10,19-hexanorandrostan-5-oic acid (**VI**)–CoA ester) (Fig. 5A3–A7; alignments of the five top models are shown in Supplementary Materials Fig. S4C) (14, 15, 27). All of these enzymes exhibited a Rossmann-like α/β/α sandwich fold characteristic of SDR-family oxidoreductases, and all were predicted to form dimers. The AlphaFold models of (3β-DH)₂, (SteA)₂, and (ScdE)₂ closely resembled those of (ScdB)₂ and (ScdG)₂, whereas the models of (3α-DH)₂ and (SteB)₂ showed different overall architectures and were more similar to each other (Fig. 5A6 and 5A7; see also alignments in Supplementary Materials Fig. S4B and S4C). Notably, models 0–2 of (SteB)₂ resembled all models of (3α-DH)₂, while models 3 and 4 of (SteB)₂ were more similar to (ScdB)₂ (Supplementary Materials Fig. S4C).

When the dimers of (3α-DH)₂ and (3β-DH)₂ were aligned, the overall complexes appeared dissimilar (Fig. 5B1). However, one monomeric subunit aligned well between the two. Aligning them as monomers revealed that 3α-DH and 3β-DH share highly similar 3D structures (Fig. 5B2). Extending this approach, we aligned all hydrogenases shown in Fig. 5A as monomers, which demonstrated that all share a conserved 3D architecture (Fig. 5B3). We subsequently aligned the amino acid sequences of these seven enzymes to identify specific regions responsible for structural divergence at the dimer level; however, no clear determinants could be identified (Supplementary Materials Fig. S4D).

ScdH, ChsH1H2, ScdD (hydratase acting on the C6–7 double bond on **IV**-CoA ester), ScdY (hydratase for the C8–14 double bond on 17-dihydroxy-9-oxo-1,2,3,4,5,6,10,19-octanorandrost-8(14)-en-7-oic acid-CoA ester), ScdN (hydratase in the β-oxidation steps following C-ring cleavage), and SteC (dehydratase for the C12β-hydroxyl group) constitute the hydratases/dehydratases identified so far in steroid degradation in TA441 (Fig. 1, 2, and Supplementary Materials Fig. S1) (28). Homology analysis indicated that ScdH, ScdY, and ScdN are members of the crotonase-like enoyl-CoA hydratase/isomerase family, whereas ScdD and ChsH1H2 belong to the MaoC-family dehydratases. SteC was suggested to be a polyketide cyclase belonging to the SnoaL/NTF2 family.

Crotonase-like hydratases typically form homohexamers (47), and AlphaFold similarly predicted hexameric assemblies for ScdH, ScdY, and ScdN (Fig. 6A1–3; expected position errors and alignments of the five top models are shown in Supplementary Materials Fig. S6A,B). Because hexameric structures are complex and difficult to compare directly, we aligned their monomers. ScdH and ScdY were highly similar, and ScdN shared local similarity on the structure around β-sheet region (Fig. 6B). Alignments of ScdH with ScdD, ChsH1, and SteC showed no detectable similarity. Conversely, alignment of ScdD with ChsH1 and ChsH2 revealed strong resemblance among ScdD, the MaoC domain of ChsH1 (ChsH1_MaoC_), and ChsH2 (Fig. 7A1–3).

**Fig. 6.**
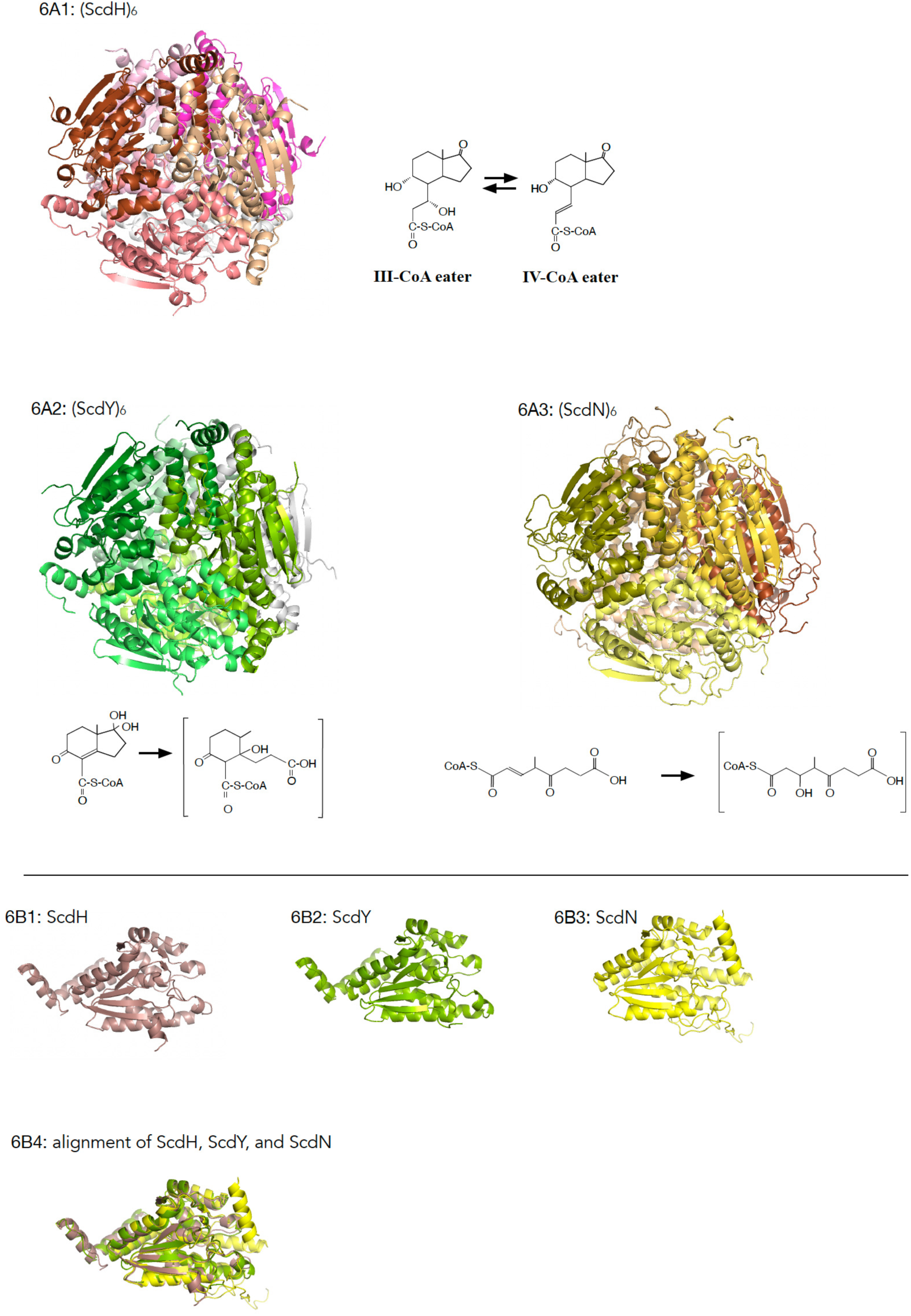
AlphaFold-predicted 3D structures of (ScdH)_6_ (6A1), (ScdY)_6_ (hydratase for the double bond at C8-14) (6A2), (ScdN)_6_ (hydratase for ³-oxidation process after C-ring cleavage) (6A3), and monomers ScdH (6B1), ScdY (6B2), ScdN (6B3). 6B4: Alignment of ScdH, ScdY, and ScdN monomers. Expected position errors and alignments of top models are shown in Supplementary Materials Fig. S6A and S6B.

**Fig. 7.**
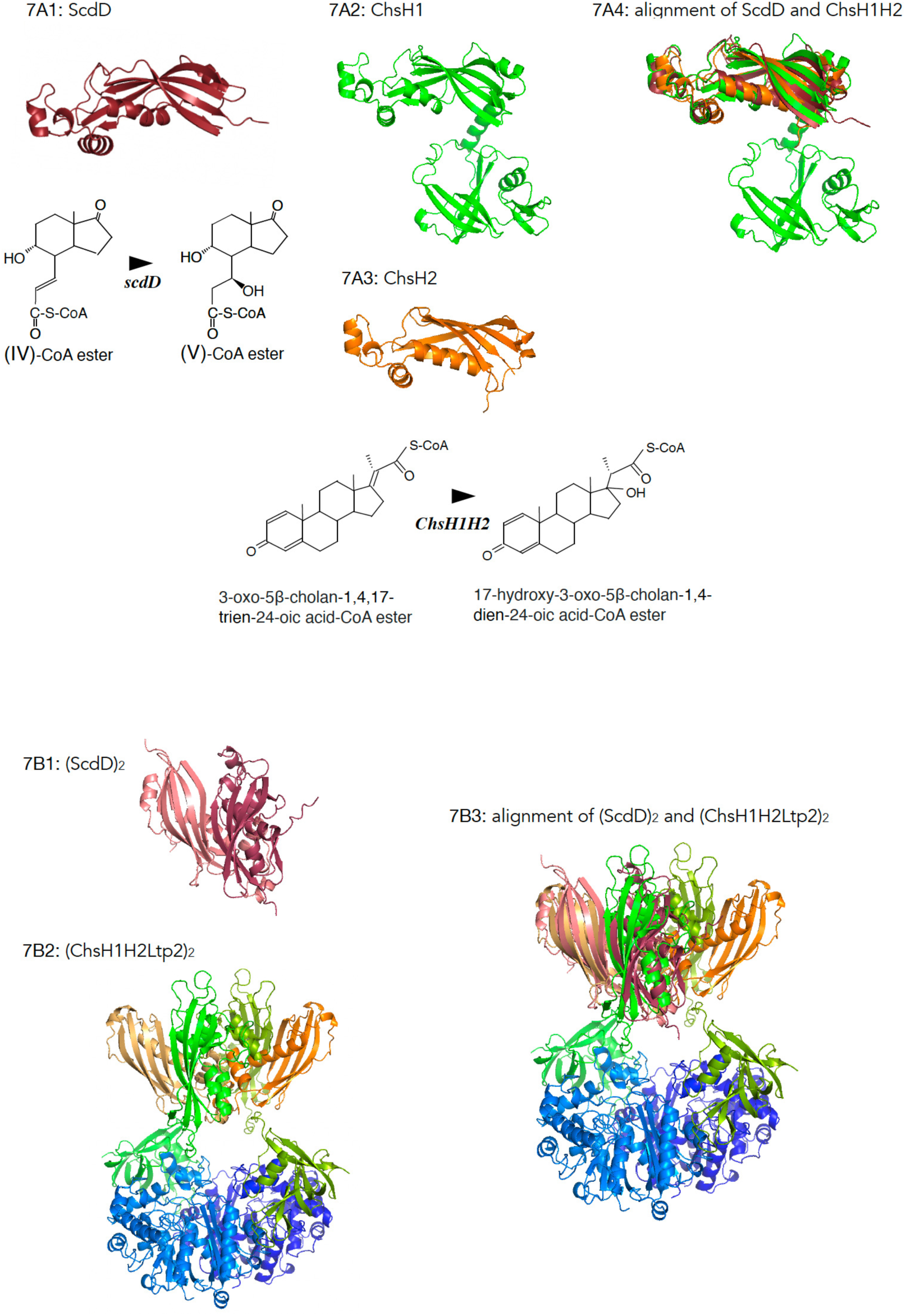
AlphaFold-predicted 3D structures of ScdD (7A1), ChsH1 (7A2), ChsH2 (7A3), (ScdD)_2_ (7B1), hetero hexamer (ChsH1H2Ltp2)_2_ (7B2), and alignment of (ScdD)_2_ and ChsH1H2Ltp2)_2_ (7B3). Reactions of ScdD and ChsH1H2 are shown below the 3D models. Expected position errors and alignments of five top models are shown in Supplementary Materials Fig. S7.

AlphaFold predicted ScdD to form a dimer (Fig. 7B1; Supplementary Materials Fig. S7), whereas previous work showed that ChsH1H2 together with Ltp2 forms a heterohexamer (ChsH1H2Ltp2)₂ in a manner similar to the *M. tuberculosis* enzyme (Fig. 7B2) (28). Superimposing the (ScdD)₂ model on (ChsH1H2Ltp2)₂ demonstrated a structural similarity between ScdD and ChsH1_MaoC_H2 (Fig. 7B3).

SteC, an NTF2-family protein, shares this classification only with ketosteroid Δ⁴-isomerase (KSI) among TA441 steroid-degrading enzymes. Monomer alignment showed similarity limited to the β-sheet region corresponding to amino acid residues ∼20–160 in SteC (Fig. 8A1–3). Bacterial NTF2-family proteins fall broadly into two groups: catalytically active dimers like KSI and catalytically inactive monomers (48). Because KSI is a dimer and AlphaFold also strongly predicted dimerization for SteC (Supplementary Materials Fig. S8A,B), we compared their dimer models; however, their overall formation was distinct (Fig. 8B1–B3). No other TA441 steroid degradation enzyme exhibited similarity to SteC. A search of other bacterial proteins identified bile acid 7α-dehydratase BaiE from *Clostridium scindens* JCM 10418/VPI 12708 (formerly *Eubacterium sp.* VPI 12708) as structurally similar (Fig. 8C1–3; Supplementary Materials Fig. S8A–C). Members of *Clostridium* are known to induce intestinal regulatory T cells (iTreg) and play major roles in mucosal immunity (49–51). BaiE is essential for converting primary to secondary bile acids (52), the latter associated with increased colorectal cancer risk (53). BaiE possesses a SnoaL-like domain within the NTF2 family (54–56). Although both are NTF2 family protein act as dehydratase on hydroxyl group on steroidal rings, SteC on C-ring and BaiE on B-ring, they share only 28% amino acid identity. Crystal analysis revealed BaiE forms a trimer (57), therefore we compare them as a trimer. Superimposition of AlphaFold-predicted trimer models of (SteC)_3_ and (BaiE) _3_ showed striking structural similarity (Fig. 8C4–6; Supplementary Materials Fig. S8D and F). Both proteins exhibited very low predicted position errors in dimer, trimer, and even tetramer models (Fig. 8C7–9; Supplementary Materials Fig. S8E and F). The reason remains unclear, but such flexibility may enable stable interactions with diverse steroid substrates.

**Fig. 8.**
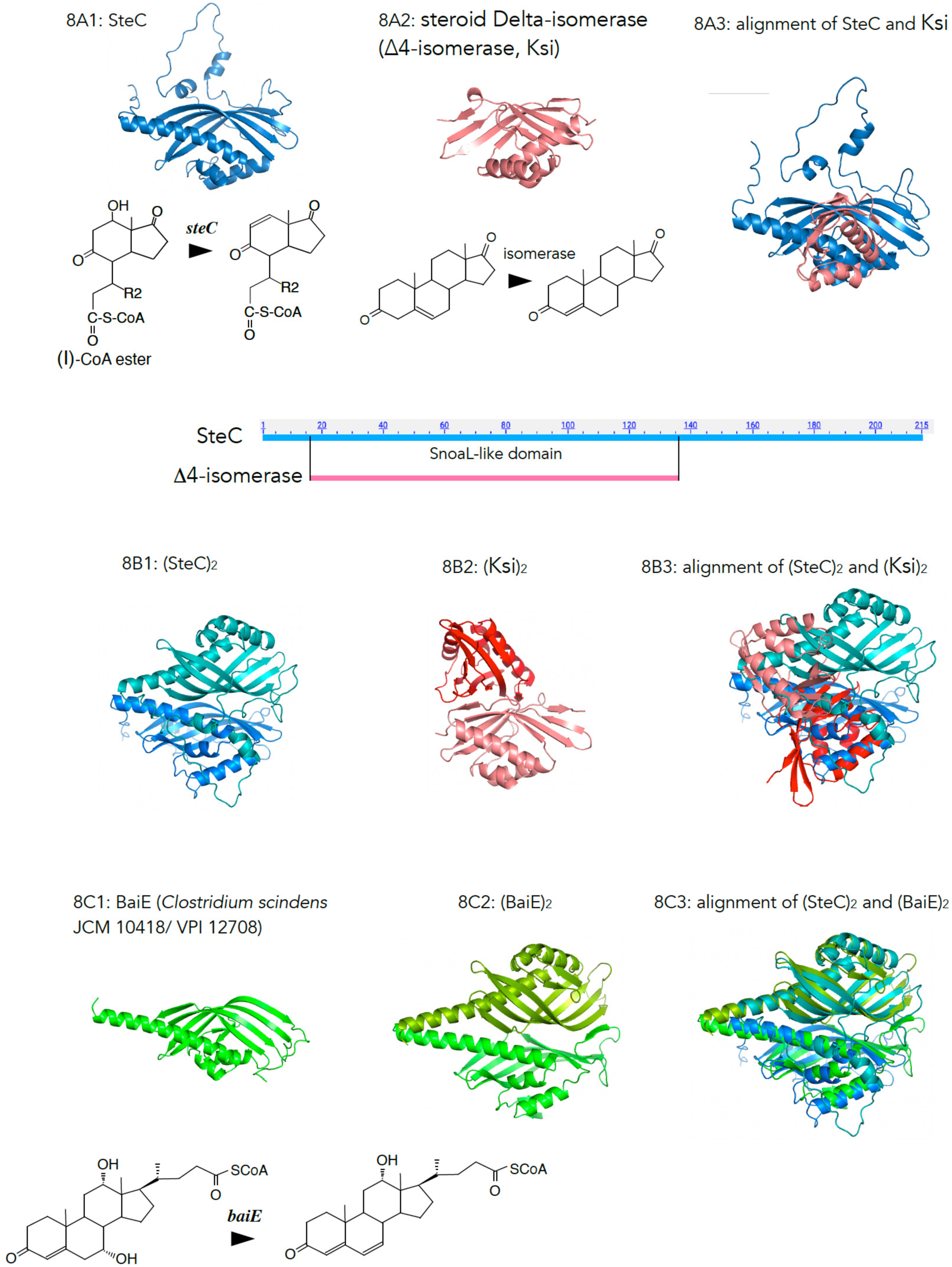
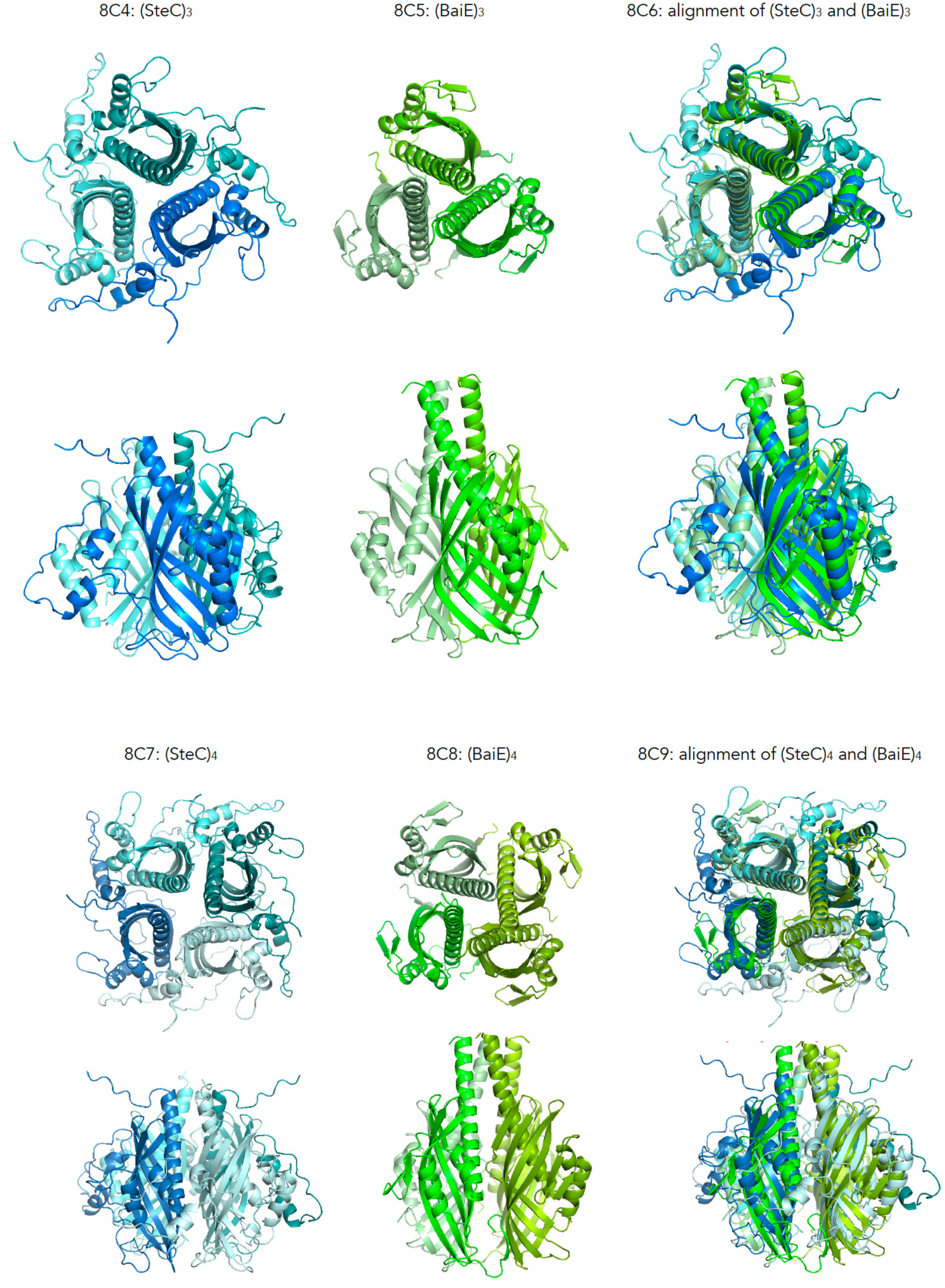
AlphaFold-predicted 3D structures of SteC (8A1), ketosteroid &4-isomerase (KSI) (8A2), alignment of SteC and KSI (8A3), dimeric structures (SteC)_2_ (7B1) and (KSI)_2_ (7B2), alignment of (SteC)_2_ and (KSI)_2_ (8B3), BaiE, bile acid 7³-dehydratase of *Clostridium scindens* JCM 10418/ VPI 12708 (8C1), (BaiE)_2_ (8C2), alignment of (SteC)_2_ and (BaiE)_2_ (8C3), trimeric structures (SteC)_3_ (8C4) and (BaiE)_3_ (8C5), alignment of (SteC)_3_ and (BaiE)_3_ (8C6), tetrameric structures (SteC)_4_ (8C7) and (BaiE)_4_ (8C8), and alignment of (SteC)_4_ and (BaiE)_4_ (8C9). Expected position errors and an alignment of five top models are shown in Supplementary Materials Fig. S8A-D.

## DISCUSSION

In this study, we identified the dehydrase responsible for converting the 7α-hydroxyl group of **III**-CoA ester to **IV**-CoA ester and the 9α-hydrogenase responsible for converting 9,17-DOHNA-CoA ester to **III**-CoA ester, which had been predicted but not previously identified, and we designated these enzymes ScdH and ScdB, respectively. As a result, all reactions occurring at the beginning of B-, C-, and D-ring degradation—specifically, those before the initiation of β-oxidation of the opened B-ring—were fully elucidated, and the degradation of the sterane structure up to D-ring cleavage, including the removal of hydroxyl groups at positions 3, 7, and 12, has now been completely clarified.

Although *scdH* and *scdB* are essential genes for B-, C-, and D-ring degradation, they are not located within the BCD-ring degradation gene cluster *tesB* through *tesR*; instead, they are encoded adjacent to *chsH2E1H1ltp2chsE2*, the gene set responsible for degradation of the isopropyl residue at C17 in C17-side chain degradation of cholic acid (ORF38 and 39 in Fig. 1).

On the opposite side of *chsH2E1H1ltp2chsE2*, 5.7 kb upstream, genes encoding 3α-dehydrogenase and KSI are located, and approximately 38 kb farther upstream lie *steABCD*, which are required for removal of the 12α-hydroxyl group, along with the BCD-ring degradation gene cluster *tesB* through *tesR* (Fig. 1). To date, the major aerobic steroid degradation pathway appears to be largely conserved among bacteria, including *M. tuberculosis* (29). Thus, elucidating steroid degradation in *Comamonas,* an opportunistic bacterium ubiquitous in the environment, is expected to provide insight into diverse bacteria–host interactions mediated by steroids.

Using AlphaFold-based structural analysis, we found that many ketone/hydroxyl-converting hydrogenases/dehydrogenases involved in steroid degradation in TA441 are SDR-family oxidoreductases that adopt Rossmann-like α/β/α folds and function predominantly as dimers. In contrast, our analysis suggested that some enzymes may adopt different three-dimensional structures depending on the stereochemistry of their substrates. These structural variations may reflect evolutionary adaptations enabling single enzymes to accommodate a wider range of steroid substrates.

Furthermore, the dehydratase SteC, which removes the C12β-hydroxyl group from derivatives of 9,17-DOHNA, was found to have a three-dimensional structure strikingly similar to that of BaiE, a bile-acid 7α-dehydratase from obligate anaerobic Gram-positive bacteria *Clostridium scindens* JCM 10418/VPI 12708. The amino-acid identity between SteC and BaiE is only ∼28%, and conventional homology searches do not detect this similarity. BaiE plays a key role in the conversion of primary to secondary bile acids by gut bacteria. The finding that *C. testosteroni* possesses a structurally analogous enzyme capable of dehydrating hydroxyl groups on the steroid nucleus may provide new clues to understanding how bacteria have acquired the ability to degrade and modify diverse steroid compounds.

## Acknowledgments

The authors sincerely thank Dr. Reizo Kato (Head of Condensed Molecular Materials Laboratory, RIKEN) and Dr. Yousoo Kim (Head of Surface and Interface Science Laboratory, RIKEN) for their thoughtful support and guidance. The authors also express their gratitude to Dr. Takemichi Nakamura, Toshihiko Nogawa, and the Support Units at RIKEN (Molecular Structure Characterization Unit, RIKEN CSRS, Wako) for their invaluable assistance with UPLC/MS analysis.

## MATERIALS AND METHODS

### Abbreviations

UPLC/MS,: ultra-performance liquid chromatography/mass spectrometry;
RT,: retention time;
MW,: molecular weight;
CoA,: coenzyme A;
3HPP,: 3-(3-hydroxyphenyl)propionic acid;
PCR,: polymerase chain reaction.

### Culture conditions

Mutant strains of *Comamonas testosteroni* TA441 were cultured at 30°C in a medium composed of equal volumes of Luria-Bertani (LB) medium and C medium, a mineral medium optimized for TA441 (9, 58). This mixed medium was used because it allows mutants to accumulate intermediate compounds more efficiently than either C medium or LB medium alone (unpublished data). Lithocholic acid (LT) was added as a filter-sterilized solution in DMSO to a final concentration of 0.05% (w/v). LT—not lithocholate—was used because C medium serves as an effective buffer and maintains the pH around 7. Similarly, 3-(3-hydroxyphenyl)propionic acid (3HPP) was prepared in acetonitrile and added to the medium at a final concentration of 0.1% (w/v). Unless otherwise indicated, cultures were incubated for 7 days for LC/MS analysis. 3HPP was added 1 day after the start of incubation, as it inhibits the growth of TA441 when added at the beginning.

### Construction of Deletion Mutants, Plasmids, and Mutants for Complementation Experiments

To construct the ORF3zntation experiments, such as pMFYMhpORF38 (Table S1), were constructed by amplifying DNA fragments containing the target ORF(s) and the Kmr gene using primers listed in Table S2. For example, primers MhpRPvuII_ORF38 and ORF38_KmrHR were used to amplify the ORF38 fragment, whereas primers ORF38_KmrH and Kmr-MFYPvuIIR were used to amplify the Kmr fragment. The amplified fragments were ligated with PvuII-digested pMFYMhpRA using the In-Fusion HD Cloning Kit.

### Ultra-High-Performance Liquid Chromatography (UPLC)/MS

A 1 mL culture was extracted twice with an equal volume of ethyl acetate under acidic conditions (adjusted to pH 2 with HCl). The ethyl acetate layer was collected, evaporated to dryness, and dissolved in 1 mL methanol. A 5 µL aliquot was injected into a UPLC/MS system (Waters Acquity UPLC H-Class–QDa; Waters, Milford, MA, USA).

### Reverse-phase liquid chromatography with tandem mass spectrometry (LC/MS/MS).

Reverse-phase LC/MS/MS analyses (Supplementary Materials Fig. S2 and S3) were performed by injecting 2 µL of samples prepared as described above for HPLC/MS. An Agilent 1100 HPLC system (Agilent, CA, USA) coupled to a 4000 QTRAP MS/MS detector (AB Sciex, Framingham, MA, USA) was used in negative ion mode with an L-column2 ODS (1.5 × 150 mm) Type L2-C18, 5 µm, 12 nm (GL Science, Tokyo, Japan). Elution was performed with 90% solution A (H₂O:formic acid = 100:0.1) and 10% acetonitrile for 1 min, followed by a linear gradient to 20% solution A and 80% acetonitrile over 7 min, maintained for 2 min. The flow rate was 0.2 mL/min. MS/MS conditions were as follows: ion source temperature, 450 °C; spray needle voltage, –4.5 kV; sheath gas pressures, 60 (gas 1) and 70 (gas 2); curtain gas, 15.

Collision energy was 20 V, and ions were detected by the Q3 detector.

## Data Availability

The authors affirm that materials and data reasonably requested by others will be made available from a publicly accessible collection or provided in a timely manner at reasonable cost and in limited quantities to members of the scientific community for noncommercial purposes.

## REFERENCES

1. Speranza A. 2010. Into the world of steroids: a biochemical "keep in touch" in plants and animals. Plant Signal Behav 5:940–943.

2. Clayton RB, Kluger RH, Augustyn A. et al. steroid. Britannica. https://www.britannica.com/science/steroid

3. Yazdan MMS, Kumar R, Leung SW. 2022. The environmental and health impacts of steroids and hormones in wastewater effluent, as well as existing removal technologies: a review. Ecologies 3:206–224.

4. Penning TM, Adamski J. 2011. Integration of steroid research: perspectives on environment factors, homeostasis in health, and disease treatment. J Steroid Biochem Mol Biol doi:10.1016/j.jsbmb.2011.04.011.

5. Dodson RM, Muir RD. 1961. Microbiological transformations. IV. The microbiological aromatization of steroids. J Am Chem Soc 83:4627–4631.

6. Gibson DT, Wang KC, Sih CJ, Whitlock H Jr. 1966. Mechanisms of steroid oxidation by microorganisms. IX. On the mechanism of ring A cleavage in the degradation of 9,10-seco steroids by microorganisms. J Biol Chem 241:551–559.

7. Sih CJ, Lee SS, Tsong YK, Wang KC. 1966. Mechanisms of steroid oxidation by microorganisms. VIII. 3,4-Dihydroxy-9,10-secoandrosta-1,3,5(10)-triene-9,17-dione. J Biol Chem 241:540–550.

8. Coulter AW, Talalay P. 1968. Studies on the microbiological degradation of steroid ring A. J Biol Chem 243:3238–3247.

9. Horinouchi M, Yamamoto T, Taguchi K, Arai H, Kudo T. 2001. Meta-cleavage enzyme gene tesB is necessary for testosterone degradation in *Comamonas testosteroni* TA441. Microbiology 147:3367–3375.

10. Horinouchi M, Hayashi T, Koshino H, Yamamoto T, Kudo T. 2003. Gene encoding the hydrolase for the meta-cleavage product in testosterone degradation by *Comamonas testosteroni*. Appl Environ Microbiol 69:2139–2152.

11. Horinouchi M, Hayashi T, Yamamoto T, Kudo T. 2003. A new bacterial steroid degradation gene cluster in *Comamonas testosteroni* TA441. Appl Environ Microbiol 69:4421–4430.

12. Horinouchi M, Hayashi T, Kudo T. 2004. Genes encoding the hydroxylase of 3-hydroxy-9,10-secoandrosta-1,3,5(10)-triene-9,17-dione in *Comamonas testosteroni* TA441. J Steroid Biochem Mol Biol 92:143–154.

13. Horinouchi M, Kurita T, Yamamoto T, Hatori E, Hayashi T, Kudo T. 2004. Steroid degradation gene cluster of *Comamonas testosteroni* consisting of 18 genes from tesB to tesR. Biochem Biophys Res Commun 324:597–604.

14. Horinouchi M, Hayashi T, Koshino H, Kurita T, Kudo T. 2005. Identification of 9,17-DOHNA, 4-hydroxy-2-oxohexanoic acid, and 2-hydroxyhexa-2,4-dienoic acid in testosterone degradation by Comamonas testosteroni TA441. Appl Environ Microbiol 71:5275–5281.

15. Horinouchi M, Kurita T, Hayashi T, Kudo T. 2010. Steroid degradation genes in *Comamonas testosteroni* TA441. J Steroid Biochem Mol Biol 122:253–263.

16. Horinouchi M, Hayashi T, Kudo T. 2012. Steroid degradation in *Comamonas testosteroni*. J Steroid Biochem Mol Biol 129:4–14.

17. Horinouchi M, Hayashi T, Koshino H, Malon M, Hirota H, Kudo T. 2014. Identification of β-oxidation products of the C-17 side chain in cholic acid degradation. J Steroid Biochem Mol Biol 143:306–322.

18. Horinouchi M, Koshino H, Malon M, Hirota H, Hayashi T. 2018. Identification of metabolites involved before D-ring cleavage. Appl Environ Microbiol 84:e01324–18.

19. Horinouchi M, Koshino H, Malon M, Hirota H, Hayashi T. 2019. Identification of octanor-seco acid derivative. J Steroid Biochem Mol Biol 185:268–276.

20. Horinouchi M, Malon M, Hirota H, Hayashi T. 2019. Identification of 4-methyl-5-oxo-octane-1,8-dioic acid in steroidal C,D-ring degradation. J Steroid Biochem Mol Biol 185:277–286.

21. Horinouchi M, Koshino H, Malon M, Hirota H, Hayashi T. 2019. Steroid degradation in *Comamonas testosteroni* TA441: entire β-oxidation cycle of the cleaved B-ring. Appl Environ Microbiol 85:e01204–19.

22. Horinouchi M, Hayashi T, Koshino H, Kudo T. 2006. ORF18-disrupted mutant accumulates 9,17-DOHNA. J Steroid Biochem Mol Biol 101:78–84.

23. Horinouchi M, Hayashi T. 2023. Comprehensive summary of steroid metabolism in TA441. Appl Environ Microbiol 89:e0014323.

24. Horinouchi M, Hayashi T. 2023. Identification of “missing links” in C- and D-ring cleavage in TA441. Appl Environ Microbiol 89:e0105023.

25. Horinouchi M, Hayashi T. 2021. Identification of CoA intermediates in C-ring degradation. Appl Environ Microbiol 87:e0110221.

26. Horinouchi M, Hayashi T, Koshino H, Malon M, Hirota H, Kudo T. 2014. Identification of 9α-hydroxy-17-oxo intermediates in *C. testosteroni* TA441. J Bacteriol 196:3598–3608.

27. Horinouchi M, Hayashi T, Koshino H, Malon M, Yamamoto T, Kudo T. 2008. Genes involved in C-12 hydroxyl inversion in cholic acid catabolism. J Bacteriol 190:5545–5554.

28. Horinouchi M. 2025. Identification of enzymes for propionyl-removal in cholic acid C-17 degradation. Microbiol Spectr 13:e0030825.

29. Horinouchi M, Hayashi T. 2025. Comprehensive review of TA441 with insights from other bacteria. Adv Appl Microbiol 131:1–20.

30. Wipperman MF, Yang M, Thomas ST, Sampson NS. 2013. FadE proteome of *Mycobacterium tuberculosis*. J Bacteriol 195:4331–4341.

31. Aggett R, Mallette E, Gilbert SE, et al. 2019. Steroid side-chain-cleaving aldolase Ltp2-ChsH2. J Biol Chem 294:11934–11943.

32. Yuan T, Yang M, Gehring K, Sampson NS. 2019. Heterohexameric hydratase–aldolase complex. Biochemistry 58:4224–4235.

33. Mohn WW, Wilbrink MH, Casabon I, et al. 2012. Cholate catabolism in *Rhodococcus* spp. J Bacteriol 194:6712–6719.

34. Crowe AM, Casabon I, Brown KL, et al. 2017. Catabolism of the last two steroid rings in *Mycobacterium tuberculosis*. mBio 8:e00321–17.

35. Holert J, Alam I, Larsen M, et al. 2014. Genome sequence of *Pseudomonas* sp. Chol1. Genome Announc 1:e000XX.

36. Feller FM, Richtsmeier P, Wege M, Philipp B. 2021. Comparative analysis of bile-salt degradation. Appl Environ Microbiol 87:e0145321.

37. Holert J, Yucel O, Jagmann N, et al. 2016. Bypass reactions in bile-salt metabolism. Environ Microbiol 18:3373–3389.

38. Wang PH, Leu YL, Ismail W, et al. 2013. Steroid core cleavage by *Steroidobacter denitrificans*. J Lipid Res 54:1493–1504.

39. Ibero J, Rivero-Buceta V, García JL, Galán B. 2022. Polyhydroxyalkanoate Production by *Caenibius tardaugens* from Steroidal Endocrine Disruptors. Microorganisms 10(4):706. doi: 10.3390/microorganisms10040706.

40. Pandey AK, Sassetti CM. 2008. Mycobacterial persistence requires host cholesterol. Proc Natl Acad Sci USA 105:4376–4380.

41. Wipperman MF, Sampson NS, Thomas ST. 2014. Cholesterol utilization by *Mycobacterium tuberculosis*. Crit Rev Biochem Mol Biol 49:269–293.

42. Ma YF, Zhang Y, Zhang JY, et al. 2009. The complete genome of *Comamonas testosteroni* reveals its genetic adaptations to changing environments. Appl Environ Microbiol 75:6812–9.

43. Yin Y, Han J, Wu H, et al. 2024. *Comamonas resistens* sp. nov. Int J Syst Evol Microbiol 74.

44. Steroid degradation – *Comamonas resistens*. KEGG. https://www.kegg.jp/kegg-bin/show_pathway?crj00984

45. Nagata Y, Nariya T, Ohtomo R, et al. 1993. Dehalogenase gene for γ-HCH degradation. J Bacteriol 175:6403–6410.

46. Jumper J, Evans R, Pritzel A, et al. 2021. AlphaFold. Nature 596:583–589.

47. Engel CK, Kiema TR, Hiltunen JK, Wierenga RK. 1998. Structure of enoyl-CoA hydratase with octanoyl-CoA. J Mol Biol 275:859–847.

48. Eberhardt RY, Chang Y, Bateman A, et al. 2013. NTF2-like superfamily. BMC Bioinformatics 14:327.

49. Ivanov II, Honda K. 2012. Intestinal commensals as immune modulators. Cell Host Microbe 12:496–508.

50. Atarashi K, Tanoue T, Shima T, et al. 2011. Colonic Treg induction by Clostridium. Science 331:337–341.

51. Atarashi K, Tanoue T, Oshima K, et al. 2013. Treg induction by selected Clostridia. Nature 500:232–236.

52. Ridlon JM, Daniel SL, Gaskins HR. 2023. The Hylemon–Björkhem pathway. J Lipid Res 64:100392.

53. Voigt W, Thomas PJ, Hsia SL. 1968. 6β-Hydroxylation of bile acids. J Biol Chem 243:3493–3499.

54. Mallonee DH, White WB, Hylemon PB. 1990. Bile acid-inducible operon from *Eubacterium*. J Bacteriol 172:7011–7019.

55. Coleman JP, White WB, Hylemon PB. 1987. Bile acid 7-dehydroxylase cloning. J Bacteriol 169:1516–1521.

56. Daniel SL, Ridlon JM. 2025. *Clostridium scindens* review. FEMS Microbiol Rev 49.

57. Bhowmik S, Chiu H-P, Jones DH, et al. 2016. Structure and function of BaiE. Proteins 84:316–331.

58. Arai H, Akahira S, Ohishi T, Maeda M, Kudo T. 1998. Phenol degradation in *Comamonas testosteroni*. Microbiology 144:2895–2903.

59. Vieira J, Messing J. 1987. Production of single-stranded plasmid DNA. Methods Enzymol 153:3–11.

60. Arai H, Yamamoto T, Ohishi T, Shimizu T, Nakata T, Kudo T. 1999. 3-(3-Hydroxyphenyl)propionic acid degradation pathway. Microbiology 145:2813–2820.

